# Technological Properties Of Milk For Cheese Making With The Use Of Antioxidants In Diets For Cows With Subtoxic Dose Of Nitrates

**DOI:** 10.1101/2020.08.02.232660

**Authors:** Marina G. Kokaeva, Rustem B. Temiraev, Magomed G. Chabaev, Roman V. Nekrasov, E.Yu. Tsis, Valery R. Kairov, Bella S. Nikkolova, Saida N. Edigova, Bela A. Konokova

**Affiliations:** North-Caucasian Mining and Metallurgical Institute (State Technological University); Science Center for Animal Husbandry named after Academy Member L.K. Ernst; Federal Science Center for Animal Husbandry named after Academy Member L.K. Ernst; Gorsky State Agrarian University; North Ossetian State University named after Kosta Levanovich Khetagurov; Maykop State Technological University

**Keywords:** cows, nitrates and nitrites, antioxidants, milk, physicochemical and technological properties, cheese making, sanitary and hygienic properties of cheese.

## Abstract

The most effective means for denitrifying products and fat oxidation processes is to prevent contamination of grain at all stages of cultivation, processing and storage. If it was not possible to prevent these processes, it is necessary to use preparations that reduce the harmful effects of nitrates, mycotoxins and peroxides. These include antioxidants, adsorbents. The aim of the research was to study the efficiency of use in diets containing a subtoxic dose of nitrates, adsorbent preparations ofSantochinum and Mold-Zap to increase the productivity, physicochemical and technological characteristics of milk of lactating cows. To achieve this goal, scientific and economic experiment was conducted on cows of the Swiss breed.Of the 40 cows, according to the method of analoguepairs, 4 groups of 10 animals each were formed. The combined use of preparations of Santochinum at a dose of 0.5 kg/t and Mold-Zap at a dose of 1.5 kg/t of compound feed contributed to an increase in the milk of cows of the 3^rd^ experimental group, as compared to the control analogues, of the fat content by 0.21%, protein - by 0.17%, density - by 0.61°A, dry matter by 0.48%, vitamin C by 48.2%, vitamin A by 51.9% with a simultaneous decrease in the nitrate content by 45.1% and nitrite - by 55, 3%. Regarding the control, the milk of the cows of the 3^rd^ experimental group had more fat globules by 11.9%, but with a smaller diameter - by 17.4%.The addition of these preparations to the diets of cows of the 3^rd^ experimental grou, contributed to an increase in protein milk by 0.21%, the proportion of casein - by 0.31%, and α-casein - by 4.56%.Inthe sample of cheese from milk of cows of the 3^rd^experimental group, there was an increase in the concentration of dry matter by 1.68%, protein in the dry matter - by 1.31%. After homogenization, in the milk of cows of the 3^rd^ experimental group, the nitrate reductase activity of xanthine oxidase exceeded 2.36 times the sample of production of cows in the control group. This technological technique contributed to the stabilization of pH=5.17 in milk, which activates the process of reducing nitrates and nitrites to ammonia. The lowest concentration of nitrates and nitrites was in the sample of cheese from the milk of animals of the 3^rd^ experimental group, exceeding in these parameters the control sample by 68.5 and 70.0%.

## Original Articles

Articles that present a contribution which is entirely new to knowledge and allow other researchers, based on the written text, to judge the conclusions, check the accuracy of the analyzes and deductions of the author and repeat the investigation if they so wish.

### 1. Relevance of the topic

The use of intensive technologies in agriculture, which implies widespread application of mineral fertilizers, is associated with the risk of accumulation of nitrates and nitrites in the soil. At the same time, factors that adversely affect photosynthesis, contribute to the accumulation of nitrates in fodder plants. Nitrates, absorbed by the leaves of plants, with the help of nitrate and nitrite reductases are restored, forming an intermediate product - hydroxylamine [1, 2, 3].**ex.** (TEMIRAEV, 2013; KOKAEVA, TEMIRAEV, BESLANEEV, CHERCHESOVA, KUBATIEVA, GUTIEVA, KOZYREV, 2017; KOKAEVA, GOGAEV, CUGKIEV, KOKAEVA, GALICHEVA, 2017.)

After being absorbed into the blood, these toxicants oxidize divalent iron of hemoglobin to ferric iron, forming methemoglobin. When poisoning with nitrates, in the blood of cows there is an increase in the level of methemoglobin, the respiratory function is impaired, milk production is decreased. In the milk containing nitrates, the dispersion of milk fat changes, the specific surface of the shell protein increases and the size of the fat globules decreases, which significantly reduces technological qualities of raw milk [4, 5, 6].

Along with this, to reduce the cost of 1 kg of milk,the producers began to make the most of thegrain of their own production in the feeding of dairy cattle. However, in the course of storage in the grain of corn, barley, wheat, etc., fat oxidation occurs with the formation of peroxides, destroying the structure of vitamins, reducing the activity of many enzymes. In addition, they are affected by mold fungi, including Aspergillus flavus and Aspergillus parasiticus, which leads to the accumulation in them of the aflatoxin B_1_ metabolite, which has a pronounced hepatotrophic effect [7, 8, 9, 10].

The most effective remedy for denitrification products, as well as the prevention of mycotoxicosis and fat oxidation processes, is to prevent contamination of grain at all stages of cultivation, processing and storage. If it was not possible to prevent these processes and it is not possible to avoid the use of this raw material as feed for dairy cattle, it is necessary to use peparations that reduce the harmful effects of nitrates, mycotoxins and peroxides. These include antioxidants, sorbents, etc. [11, 12, 13].

**The aim of the research** was to study the effectiveness of use in the diets containing a subtoxic dose of nitrates, preparations of Santochinum (with antioxidant properties) and Mold-Zap (mold inhibitor), to improve the productivity, physical, chemical and technological characteristics of milk of lactating cows.

## Material and research methods

To achieve the goal, on cows of the Swiss breed in the conditions of the farm “Meat Products” of the RNO-Alania, scientific and economic experiment was conducted. Of the 40 cows, selected on the base of breed, age in calving, live weight, date of last calving, previous lactation productivity and fat content in milk, 4 groups of 10 animals each were formed according to the method of analogue pairs.

In the course of the research, the cows in the control group received the basal diet (BD), and to the diets of animals of the 1, 2, and 3 experimental groups, test preparations were added to the BD at doses determined by the scheme of the experiment (Table 1).

**Table 1.**
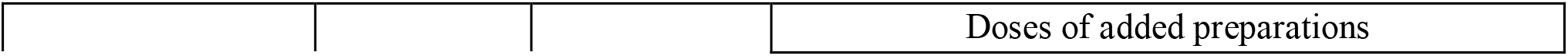
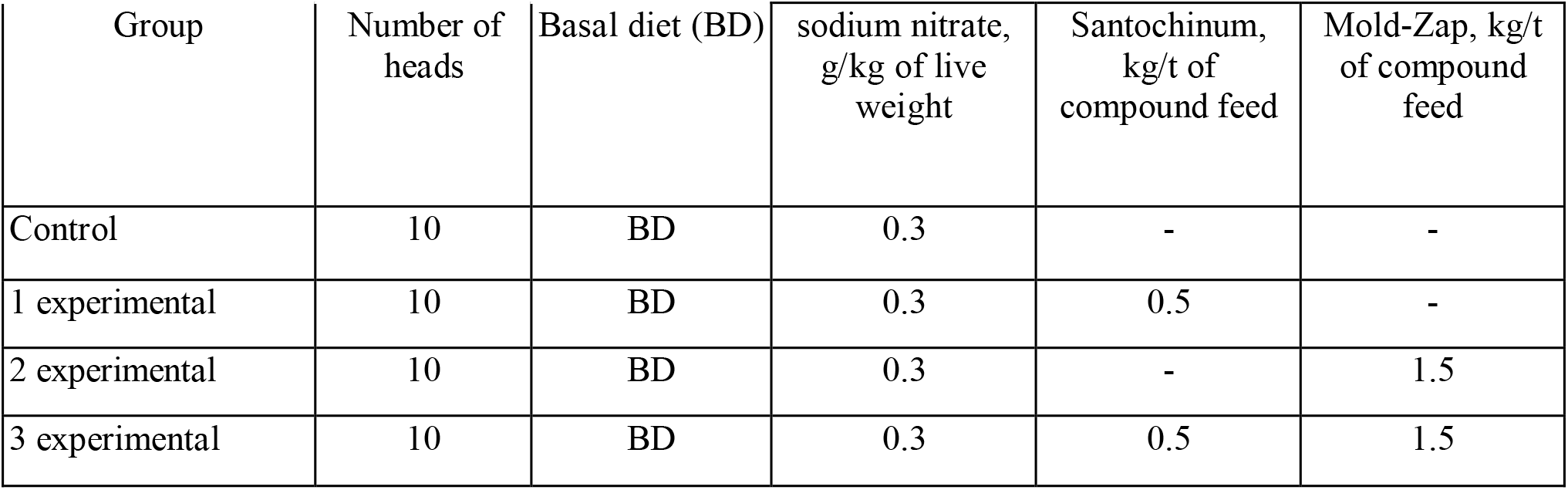
Scheme of the scientific and economic experiment.

Feeding the experimental cows was carried out with rations balanced in accordance with detailed standards of the RAAS. When preparing the rations, the sugar-protein ratio was strictly observed due to feeding of the molasses. In regularly selected medium samples of feed and their residues, the chemical composition and nutritional value were determined according to the generally accepted method [14].

Milk productivity of cows was determined by the results of the control milk yield, carried out once a decade. On the same days, the fat content in milk was determined by the Gerber acid method. The protein content in milk was determined by the formol method.

To assess the technological properties of the milk of cows of all groups, the samples of Ossetian pickled cheese were produced in accordance with GOST 4991-81 “Pickled Cheese”. The product is produced from the volume of milk yield in 24 hours. The adequacy of the milk of experimental cows for making cheese was determined by the time of rennet digestion of milk, while the acidity and temperature of the milk were recorded.

The concentration of nitrates and nitrites in the blood, milk and cheese was determined by the method described by Skorodinsky, Z.P. (1977) [15].

All the results obtained in the course of the research were statistically processed by the Student’s criterion.

## Research results and discussion

In the feeds that were used in the summer and winter diets of experimental animals, we studied the content of nitrate and nitrite ions and aflatoxin B_1_ (Table 2).

**Table 2.** Content of nitrates, nitrites and aflatoxin B_1_ in feed, mg / kg.

| Kopma | NO<sub>3</sub><sup>-</sup> |  | NO<sub>2</sub><sup>-</sup> |  | Aflatoxin B<sub>1</sub> |  |
| --- | --- | --- | --- | --- | --- | --- |
|  | MPC | actual | MPC | actual | MPC | actual |
| Oat-pea grass | 500 | 14 | 10 | 0.31 | - | - |
| Rape grass | 500 | 11 | 10 | 0.25 | - | - |
| Grass of artificial pasture | 500 | 14 | 10 | 0.11 | - | - |
| Cereal and mixed grass hay | 1000 | 44 | 10 | 0.25 | 0.05 | traces |
| Corn silage | 500 | 12 | 10 | 9.13 | 0.05 | traces |
| Compound feed | 300 | 3 | 10 | 0.03 | 0.05 | 0.03 |
| Feed molasses | 1500 | 27 | 10 | 0.34 | - | - |
| Fodder beet | 2000 | 36 | 10 | 0.27 | - | - |

None of the feeds that were included in the rations of experimental animals exceeded the maximum permissible concentrations (MPC) for the content of nitrate and nitrite ions, as well as aflatoxin B_1_. This testifies to the self-purification of the soil of the farm from nitrates and nitrites, since during the last 12-15 years nitrogen fertilizers have almost not been applied to feed crops because of their high cost. On this basis, to evaluate the denitrification properties of the tested preparations in the rations of experimental animals, taking into account the content of nitrate ions in the feed, included sodium nitrate, so that the level of nitrates in them was subtoxic - not more than 0.03 g / kg of body weight of cows [16].

The introduction of the tested preparations into the rations of animals of the compared groups had practically no effect on their palatability.

According to the results of the control milk yields, the milk productivity of the experimental animals and the feed consumption per unit of production were determined (Table 3).

**Table 3.**
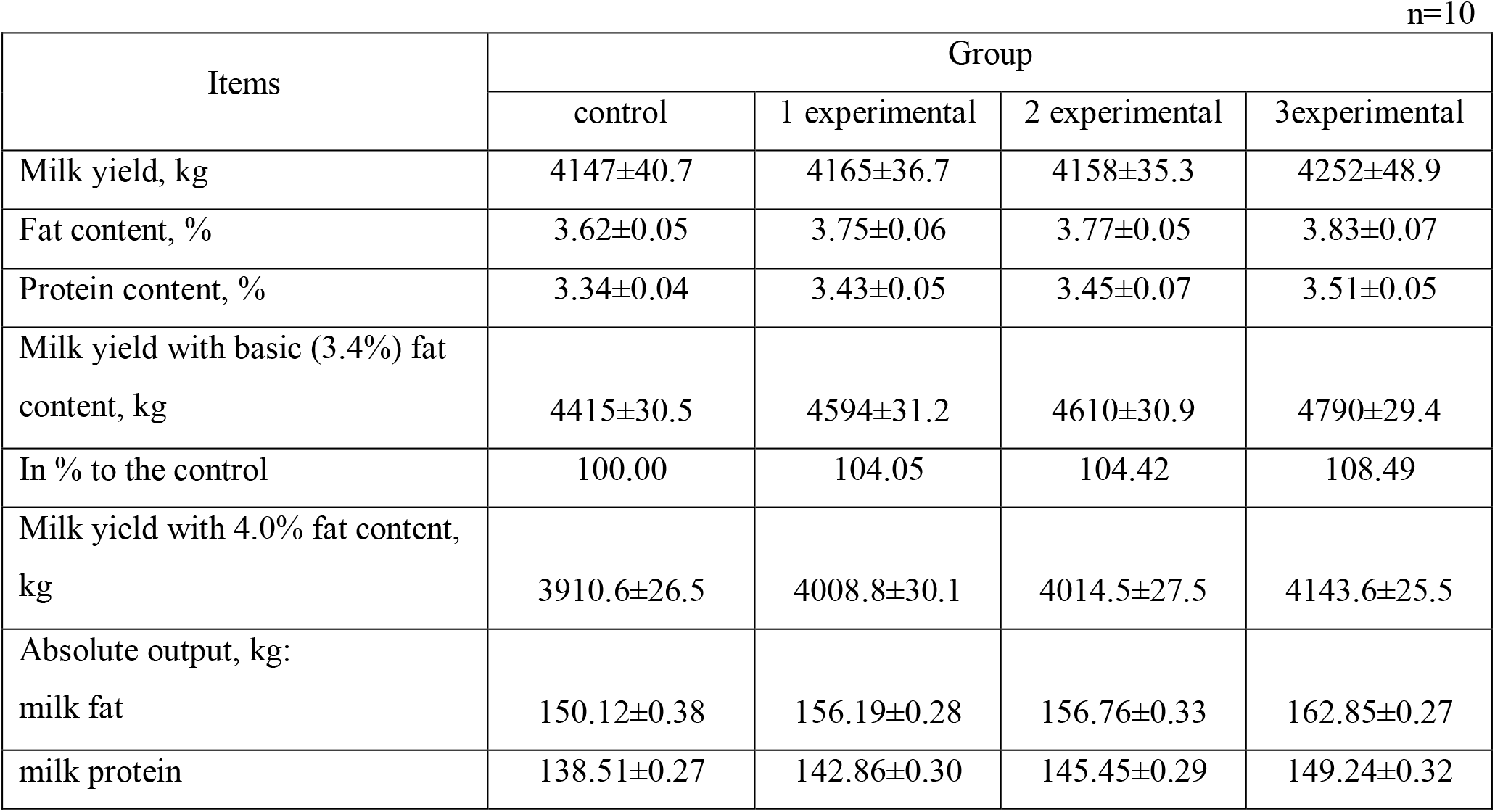
Dairy productivity of cows and feed consumption per unit of production.

| Items | Group |  |  |  |
| --- | --- | --- | --- | --- |
|  | control | 1 experimental | 2 experimental | 3experimental |
| Milk yield, kg | 4147±40.7 | 4165±36.7 | 4158±35.3 | 4252±48.9 |
| Fat content, % | 3.62±0.05 | 3.75±0.06 | 3.77±0.05 | 3.83±0.07 |
| Protein content, % | 3.34±0.04 | 3.43±0.05 | 3.45±0.07 | 3.51±0.05 |
| Milk yield with basic (3.4%) fat content, kg | 4415±30.5 | 4594±31.2 | 4610±30.9 | 4790±29.4 |
| In % to the control | 100.00 | 104.05 | 104.42 | 108.49 |
| Milk yield with 4.0% fat content, kg | 3910.6±26.5 | 4008.8±30.1 | 4014.5±27.5 | 4143.6±25.5 |
| Absolute output, kg: |  |  |  |  |
| milk fat | 150.12±0.38 | 156.19±0.28 | 156.76±0.33 | 162.85±0.27 |
| milk protein | 138.51±0.27 | 142.86±0.30 | 145.45±0.29 | 149.24±0.32 |

When comparing the actual yield, it can be seen that cows from the control group produced 105 kg or 2.5% less milk for lactation compared to the 3^rd^ experimental group, but according to statistical processing the difference was not significant (P>0.05).

In the course of the research, the fat content in the milk of cows in the control group averaged 3.62% per lactation. This indicator was higher in the milk of cows of the 3^rd^ experimental group - 3.83%, which is 0.21% more than in the control (P <0.05).

The combined use of preparations Santochinum and Mol-Zap in the diets of dairy cattle with a subtoxic dose of nitrates contributed to an increase in the protein level in milk, thanks to this, in the milk of cows of the 3^rd^experimental group, the protein content was 0.17% (P<0.05) higher than in the control.

When detoxifying nitrates using a mixture of these preparations in the mammary gland of lactating cows, the synthesis of milk fat and protein is optimized, as evidenced by a significant (P<0.05) increase in the absolute yield of milk fat and protein in animals of the 3^rd^experimental group versus the control analogues by 12.73 kg or 8.5% and by 10.73 kg or 7.7% respectively.

For greater objectivity, milk yield of natural fat content was converted into milk of 3.4% (basic) and 4% fat content. According to these indicators, cows of the 3^rd^ experimental group also had the best results - 4,415 and 4,143.6 kg, which is by 8.49% and 5.90% significantly (P<0.05) more than the cows of the control group.

To study the effectiveness of the use of the tested feed preparations, we studied some physicochemical properties of the milk of experimental cows (Table 4).

**Table 4.** Physical and chemical properties of milk of experimental cows.

| Item | Group |  |  |  |
| --- | --- | --- | --- | --- |
|  | control | 1 experimental | 2 experimental | 3 experimental |
| Density, A° | 27.72±0.10 | 28.12±0.12 | 28.18±0.13 | 28.33±0.14 |
| Acidity, T° | 18.21±0.12 | 17.94±0.13 | 17.84±0.12 | 17.70±0.15 |
| Dry matter, % | 12.30±0.14 | 12.63±0.12 | 12.68±0.11 | 12.78±0.15 |
| Milk fat, % | 3.62±0.05 | 3.75±0.06 | 3.77±0.05 | 3.83±0.07 |
| Somatic cells, % | 8.68±0.04 | 8.88±0.05 | 8.91±0.04 | 8.95±0.06 |
| Milk protein, % | 3.34±0.04 | 3.43±0.05 | 3.45±0.06 | 3.51±0.05 |
| Lactose, % | 4.51±0.05 | 4.58±0.08 | 5.46±0.05 | 4.61±0.02 |
| Ash, % | 0.83±0.005 | 0.87±0.004 | 0.85±0.004 | 0.83±0.005 |
| Vitamin C, mg/l | 13.79±0.22 | 17.33±0.30 | 17.86±0.25 | 20.44±0.27 |
| Vitamin A, mg/l | 0.268±0.002 | 0.363±0.004 | 0.377±0.004 | 0.407±0.005 |
| Ammonia, mg/l | 2.08±0.05 | 2.63±0.07 | 2.73±0.07 | 3.14±0.09 |
| Nitrates, mg/l | 7.76±0.13 | 5.96±0.18 | 5.75±0.21 | 4.26±0.18 |
| Nitrites, mg/l | 0.152±0.004 | 0.102±0.003 | 0.097±0.005 | 0.068±0.006 |
| AflatoxinM<sub>1</sub>), mg/kg<br>(MPC=0.005 mg/kg) | 0.0047<br>±0.0003 | 0.0037<br>±0.0002 | 0.0034<br>±0.0004 | 0.0024<br>±0.0007 |

In the milk of animals of the control group, the acidity index was 18.21°T, which is 0.51°T (P<0.05) lower compared to the production of cows of the 3^rd^experimental group, that is, with joint addition of preparations, the negative effect of nitrates and nitrites on the analyzed indicator decreases.

The density of milk of animals is directly dependent on the concentration of dry matter. In the milk of cows of the control group, the dry matter content was 12.30%, and its density was 27.72°A. The joint addition of preparations Santochinum and Mold-Zap had a positive effect on these indicators of the milk of animals from the 3^rd^ experimental group, which allowed them to reliably (P<0.05) exceed the control in density by 0.61°A and the concentration of dry matter in products - by 0.48%. Moreover, these indicators of milk of cows of all groups were within the normal range.

Due to the high antioxidant properties, the preparations Santochinum and Mold-Zap had a positive effect on metabolismand growth of vitamin-synthesizing bacteria Flavobacteriumvitaramen in the rumen, therefore the milk of cows of the 3^rd^experimental group compared to the control analogues turned out to be significantly (P<0.05) more saturated with vitamin C by 48.2% and vitamin A - by 51.9%.

Nitrate- and nitrite reductase of the microflora of the rumen restore nitrates and nitrites to ammonia, which is used by other protozoa for the synthesis of protein of their own body. Therefore, in the course of research, an inverse relationship was established between the concentration of nitrates and nitrites in milk, on the one hand, and ammonia, on the other. On this basis, the highest ammonia content was in the milk of cows of the 3^rd^experimental group - 3.14 mg/l, which is 50.9% (P<0.05) more than in the control.

Joint addition of the tested preparations provided the highest degree of denitrification of cows’ products from the 3^rd^experimental group, due to which their milk had a significantly (P<0.05) lower nitrate content by 45.1% and nitrites - by 55.3% than in the control.

Along with this, the milk of animalsof the 3^rd^experimental group contained significantly less aflatoxin M_1_ (metabolite of aflatoxin B_1_) by 48.9% than in the control. The content of this mycotoxin in the milk of cows of the compared groups was below the MPC.

Consequently, the enrichment of rations of lactating cows with a high content of nitrates with a mixture of preparations Santochinum and Mold-Zap had a positive impact on the physicochemical and sanitary and hygienic properties of milk.

It is known that it depends on the content of milk protein and fat for what purposes dairy raw materials will be used: for cheese-making or for butter-making. Given the superiority of the cows of the experimental groups over the control analogues in terms of the fat content in milk, we studied the constants, diameter and the number of fat globules in their milk (Table 5).

**Table 5.** Constants, diameter and number of fat globules of experimental cows.

| Item | Group |  |  |  |
| --- | --- | --- | --- | --- |
|  | control | 1 experimental | 2 experimental | 3 experimental |
| Diameter of fat globules, $\mu\text{m}$ | $3.38 \pm 0.10$ | $2.90 \pm 0.16$ | $3.00 \pm 0.11$ | $2.79 \pm 0.11$ |
| Number of fat globules, billion / $\text{cm}^3$ | $5.69 \pm 0.10$ | $6.05 \pm 0.11$ | $5.97 \pm 0.17$ | $6.37 \pm 0.11$ |
| Iodine number | $33.48 \pm 0.18$ | $32.50 \pm 0.18$ | $32.61 \pm 0.19$ | $31.59 \pm 0.19$ |
| Peroxide value | $0.053 \pm 0.002$ | $0.062 \pm 0.002$ | $0.058 \pm 0.002$ | $0.074 \pm 0.002$ |
| Fat content in milk, % | $3.62 \pm 0.05$ | $3.75 \pm 0.06$ | $3.77 \pm 0.05$ | $3.83 \pm 0.07$ |

The research results showed that when using preparations Santochinum and Mold-Zap in combination, there was a decrease in the oil-producing properties of the milk. As compared to the control, in the milk of the cows of the 3^rd^experimental group there were significantly (P<0.05) more fat globules by 11.9%, but with a smaller diameter - by 17.4% (P<0.05) with simultaneous deterioration of the constants (iodine and peroxide value) of dairy raw materials.

On this basis, we decided to evaluate the cheese-making qualities of the milk of experimental cows, for which the content of fractions and the diameter of casein micelles were determined (Table 6).

**Table 6.** The content of fractions and the diameter of casein micelles.

| Item | Group |  |  |  |
| --- | --- | --- | --- | --- |
|  | control | 1 experimental | 2 experimental | 3 experimental |
| Average protein content in milk, % | 3.34±0.04 | 3.43±0.05 | 3.45±0.06 | 3.51±0.05 |
| Casein portion, % | 2.40±0.02 | 2.58±0.03 | 2.62±0.02 | 2.71±0.04 |
| Casein content, %: |  |  |  |  |
| α-casein | 31.57±0.21 | 35.16±0.22 | 34.61±0.18 | 36.13±0.26 |
| β-casein | 53.41±0.24 | 53.71±0.30 | 53.55±0.28 | 53.62±0.33 |
| γ-casein | 15.02±0.19 | 11.13±0.16 | 11.84±0.19 | 10.25±0.13 |
| Total | 100.0 | 100.0 | 100.0 | 100.0 |
| Casein micelle diameter, °A | 610±3.2 | 663±3.0 | 675±4.0 | 711±3.5 |

Adding a mixture of these preparations to the diets of cowsof the 3^rd^ experimental group contributed, against the control, to a significant (P<0.05) increase in milk of the milk protein by 0.21% and the proportion of casein in it - by 0.31%.

Considering that from the casein fractions onlyα- and β-casein fractions coagulate under the action of rennet, it was found that the use of preparations Santochinum and Mold-Zap as acomplex did not affect the concentration of β-casein in milk, but provided a reliable (P<0.05) an increase in the proportion of α-casein in the production of analogues of the 3^rd^ experimental group by 4.56% while reducing the content of the γ-fraction of casein - by 4.77 (P<0.05) than in the control.

In the course of our research, it was confirmed that in the process of denitrification with increasing levels of α-casein, there is an increase in the diameter of casein micelles in the production. With this in mind, relative to the control analogues, the diameter of the micelles of casein milk of animals of the 3^rd^experimental group was significantly (P<0.05) greater by 16.6%.

The milk fat globules contain the enzyme xanthine oxidase, which, when the pH is close to 5.15, exhibits nitrate reductase activity, which makes it possible to reduce the concentration of nitrates and nitrites in dairy products. In the process of milk homogenization, the milk fat globule membranebreaks and releases this enzyme. Therefore, before introduction of the rennet, we subjected normalized raw milk to homogenization.

Taking into account the higher stimulating effect of these preparations on the protein content in milk, it was important to study the possibility of processing the milk of animals of the compared groups on samples of Ossetian cheese (Table 7).

**Table 7.**
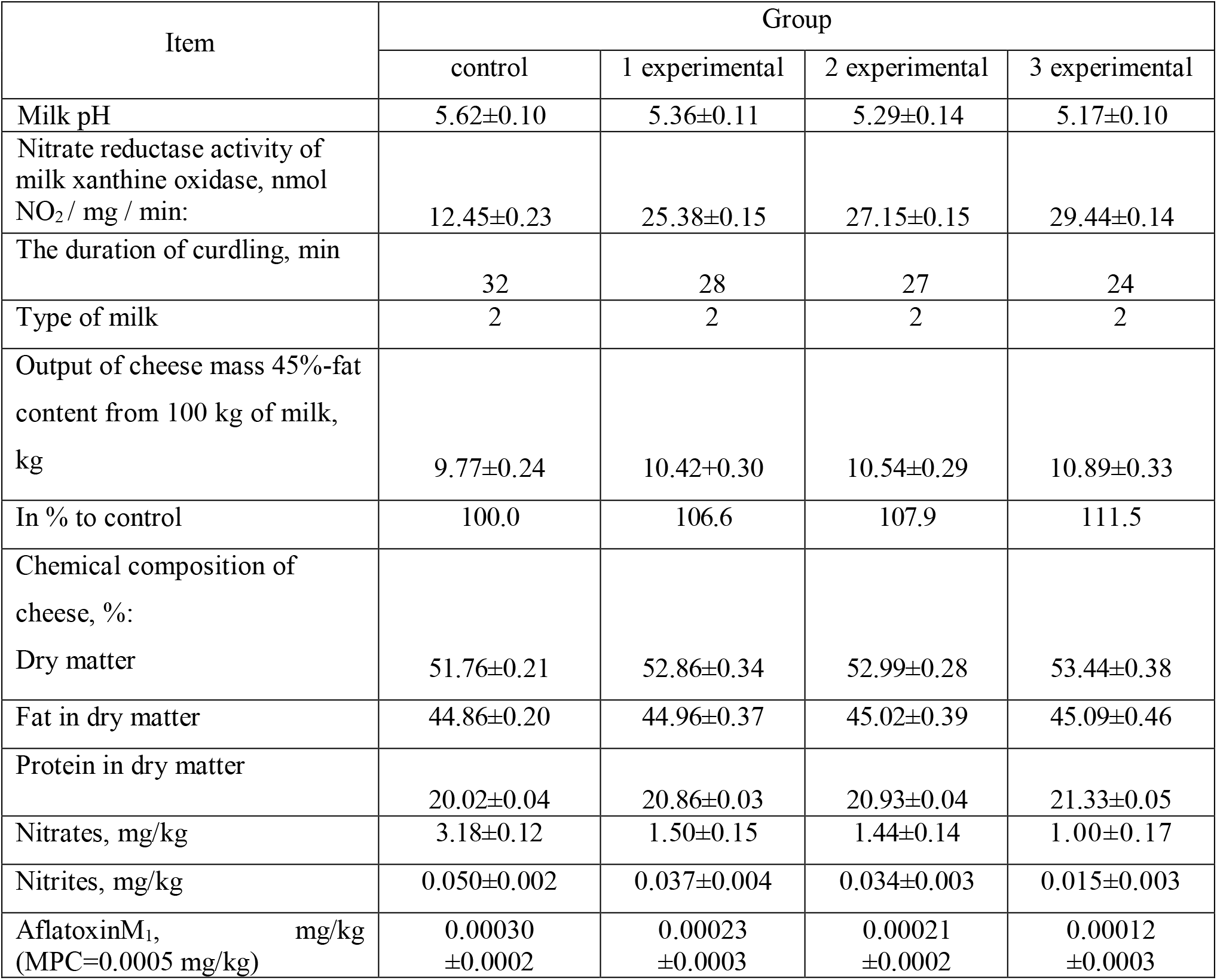
Cheese-making properties of milk of experimental cows.

It has been found that according to the adequacy for cheese making, the milk from cows of the compared groups corresponded to the second (desirable) type. However, if the milk of animals of the control group was curdled for 32.0 minutes, then the milk of cows of the 3^rd^experimental group - 8 minutes faster.

Improvement of protein metabolism under the action of a mixture of antioxidant preparations allowed cows of the 3^rd^experimental group to ensure the highest yield of cheese mass of 45% fat content - 10.89 kg, which is significantly (P<0.05) more than in the control - by 11.5%. At the same time, the rennet clot of all samples was dense and elastic with normal syneresis.

In the course of the research, the joint addition of preparations Santochinum and Mold-Zap had a more favorable effect on the chemical composition of the cheese. Therefore, in the sample of cheese from the milk of cows of the 3^rd^ experimental group, there was a significant (P <0.05) increase in the dry matter concentration by 1.68% and in protein in the dry matter - by 1.31% as compared to the control sample.

After homogenization in the milk of cows of the 3^rd^experimental group, the nitrate reductase activity of xanthine oxidase was intensified 2.36 times against the sample of production of cows in the control group. This technological method, along with an increase in the nitrate reductase activity of xanthine oxidase, helped stabilize the pH = 5.17 of milk, which activates the process of reducing nitrates and nitrites.

On this basis, the rennet clot obtained from the milk of animals of the3^rd^ experimental group differed by the lowest concentration of nitrates and nitrites, significantly (P <0.05) exceeding in these parameters the cheese mixture of the control sample, respectively, by 68.5 and 70.0%.

It should be noted that in the sample of cheese from the milk of cows of the 3^rd^experimental group, the content of aflatoxin M_1_ was 0.00012 mg / kg, which is 60.0% (P <0.05) less than in the control. The content of this mycotoxin in samples of cheese from milk of cows of the compared groups was below the MPC.

## Conclusions

1. Joint inclusion in the rations with a subtoxic dose of nitrates antioxidants Santochinum in a dose of 0.5 kg/t and Mold-Zap in a dose of 1.5 kg/t of compound feed in lactating cows of the 3^rd^experimental group provided reliable (P<0. 05) increase in milk fat content by 0.21% and protein - by 0.17%, milk yield indicator of(basic) fat content 3.4% - by 375 kg or 8.49%, as well as an increase in the absolute yield of milk fat - by 8.5 and protein - by 7.7%, respectively, as compared to the control.
2. The combined use of preparations Santochinum and Mold-Zap for denitrification contributed in cows of the 3^rd^experimental group, against the control, to the improvement of the physicochemical indicators of milkwhich resulted in:

- areliable (P<0.05) density superiority by 0.61°A and dry matter concentration - by 0.48%;
- significantly (P<0.05) higher saturation with vitamin C by 48.2% and vitamin A - by 51.9%;
- an increase in the ammonia content by 50.9% (P<0.05) with a simultaneous significant (P<0.05) decrease in the level of nitrates - by 45.1% and nitrites - by 55.3%;
- significantly (P<0.05) lower content of aflatoxin M_1_ by 48.9%, while its level in the milk of cows of the compared groups was lower than the MPC.
3. Animalsof the 3^rd^ experimental group, the rations of which included a mixture of the tested preparations, were characterized by the best technological properties of milk, which against the control analogues was expressed:

- in a significant (P<0.05) increase in the content of casein by 0.31%, the proportion of α-casein in it - by 4.56% with a simultaneous increase in the diameter of casein micelles - by 16.6% (P<0.05);
- in a significant (P<0.05) increase in the output of the cheese mass of 45% fat content - by 11.5%;
- in a significant (P<0.05) increase in the cheese sample of the dry matter concentration by 1.68%, protein in the dry matter - by 1.31%;
- in a reliable (P<0.05) decrease in the concentration of nitrates and nitrites in the cheese sample by 68.5 and 70.0%, respectively;
- in a reliable (P<0.05) decrease in the content of aflatoxin M_1_ by 60.0% (P<0.05), while the content of this mycotoxin in samples of cheese from cow’s milk of the compared groups was below the MPC.

